# “Global Transcriptional Response to CRISPR/CAS9-AAV6 Based Genome Editing” Matches Transcriptional Response to Specific Small Molecule Perturbations

**DOI:** 10.1101/399311

**Authors:** Akpéli V. Nordor, Martin J. Aryee, Geoffrey H. Siwo

## Abstract

Cromer *et al.* [1] recently reported global transcriptional changes occuring in cells in response to CRISPR/Cas9 gene editing. Using a CD34^+^ hematopoietic and progenitor stem cell model, they observed differentially expressed genes enriched for immune, stress and apoptotic processes following treatment with a CRISPR/Cas9-AAV6 genome-editing system. Following treatment with Cas9’s mRNA they observed transcriptional changes enriched for viral response as well as a downregulation of metabolic and cell cycle processes. Similarly, they observed a downregulation of metabolic processes in response to electroporation. Surprisingly, no enrichment for viral response genes was observed following treatment with AAV6 while minor transcriptional changes enriched for DNA damage signature occurred in response to Cas9/sgRNA ribonucleoprotein.

---

We hypothesized that beyond enrichment for various key cellular pathways, transcriptional response associated with CRISPR/Cas9 gene editing machinery could potentially bear similarities to transcriptional responses associated with specific small molecules or drugs. Such shared transcriptional responses could be relevant in predicting small molecule effects on genome editing, in identifying small molecules that could be used to modulate diffferent aspects of genome editing, and in predicting potential pharmacological interactions between genome editing technologies and small molecules. To test our hypothesis, we used the next generation Connectivity Map (CMap), a large-scale compendium of functional perturbations in cancer cell lines and their corresponding gene expression readouts.

This resource facilitates the discovery of connections between genes, drugs, and diseases, and is accessible through an online software platform (available at https://clue.io) [2, 3]. Briefly, the CMap pattern-matching algorithm scores the similarity between a query gene expression signature and the CMap perturbagen reference signatures (signatures of small molecule compounds here). The enrichment score computed by CMap is referred to as a “connectivity score”. A positive (negative) connectivity score indicates that the perturbation induced by the treatment corresponding to the query signature goes in the same (opposite) direction as the perturbation induced by the CMap compound. The higher the connectivity score is, the higher is the similarity between the query and reference signature (CMap compound). We constructed the query signatures by extracting the lists of up-regulated and down-regulated genes from the supplementary information reported by Cromer *et al.* (Supplementary Table S1) for each condition tested: electroporation of Cas9 mRNA and sgRNA, “mRNA”; electroporation of Cas9 mRNA and sgRNA followed by transduction of AAV6 vector, “mRNA + AAV”; electroporation of Cas9 protein complexed with sgRNA, “RNP”; electroporation of Cas9 protein complexed with sgRNA followed by AAV6 transduction, “RNP + AAV”; mock electroporation “Elect.”; and AAV6 transduction alone, “AAV”. The sgRNA in each case was designed to target the hemoglobin B (*HBB*) locus which is a clinically relevant model for sickle cell disease (in the scope of several translational research programs leveraging CRISPR/Cas9 technology). All experiments in Cromer *et al* [1] were performed in CD34^+^ hematopoietic stem cells obtained from four individual donors to account for potential individual variability to genome editing machinery [4]. The query signature lists were restricted to genes belonging to 10,174 best inferred genes (BING) in the L1000 assay. We used the significance threshold MLogP ≥ 7 consistently with the Cromer *et al.* original report and limited sizes of gene lists to 150 in accordance to the technical limit of the CMap (R code is available on GitHub: https://github.com/siworesearch/crispr_cmap).

We found that transcriptional responses to several small molecule perturbations and components of CRISPR/Cas9-AAV6 editing machinery show close similarities. While the connectivity score suggested strong positive connectivity to CRISPR/Cas9-AAV6 in some cases (Figure 1), it also indicated strong negative connectivity in other cases (Figure 2). Considering the top 20 small molecules with a high positive connectivity score suggesting similarity in global transcriptional responses to Cas9 mRNA, four – etoposide, teniposide, amsacrine and idarubicin – are topoisomerase inhibitors (Figure 1A and Supplementary Table S1). Notably, gene expression response to other CRISPR/Cas9 editing components, i.e. mRNA + AAV, RNP, RNP + AAV, Elect., AAV, was also suggested to be directly similar to gene response to topoisomerase inhibitors in the top 20 small molecules (Figure 1B to 1F). Topoisomerase inhibitors identified for the other gene editing components are etoposide (mRNA + AAV, RNP, RNP + AAV and AAV), teniposide (mRNA + AAV, RNP + AAV and AAV), amsacrine (mRNA + AAV and AAV), idarubicin (mRNA + AAV, RNP, RNP + AAV and AAV), amonafide (RNP), temozolomide (Elect.), and irinotecan (AAV). It is important to note that while these results suggest shared transcriptional signatures between each of the CRISPR/Cas9 gene editing components and topoisomerase inhibitors, the fact that some of the topoisomerase inhibitors share similarities to only a subset of the editing components suggests distinct underlying molecular mechanisms. The shared transcriptional responses with topoisomerase inhibitors is consistent with the critical role of DNA repair pathways in gene editing [5–7]. Furthermore, besides the topoisomerase inhibitors, we identified other DNA repair-targeting small molecules with positive connectivity to the CRISPR/Cas9 genome editing components. These include anisomycin (mRNA and mRNA + AAV), chlorambucil (RNP and RNP + AAV), olaparib (RNP + AAV), serdemetan/ JNJ-26854165 (RNP + AAV), clofarabine (RNP + AAV and AAV), rucaparib (Elect.), AG-14361 (Elect.) and cladribine (AAV). Interestingly, one of these compounds – serdemetan – inhibits MDM2, a negative regulator of P53 [8, 9]. Recently [10], CRISPR/Cas9 was shown to induce a P53-mediated DNA damage response and that P53 inhibits CRISPR-Cas9 gene editing [11]. Our results showing shared transcriptional responses between serdemetan and RNP + AAV supports the role of P53-mediated DNA damage response in CRISPR/Cas9 editing. We provide additional information for other top 20 small molecules including various kinases and protein synthesis inhibitors with positive connectivity to the various genome editing components in Figure 1 and Supplementary Table S1.

**Figure 1:**
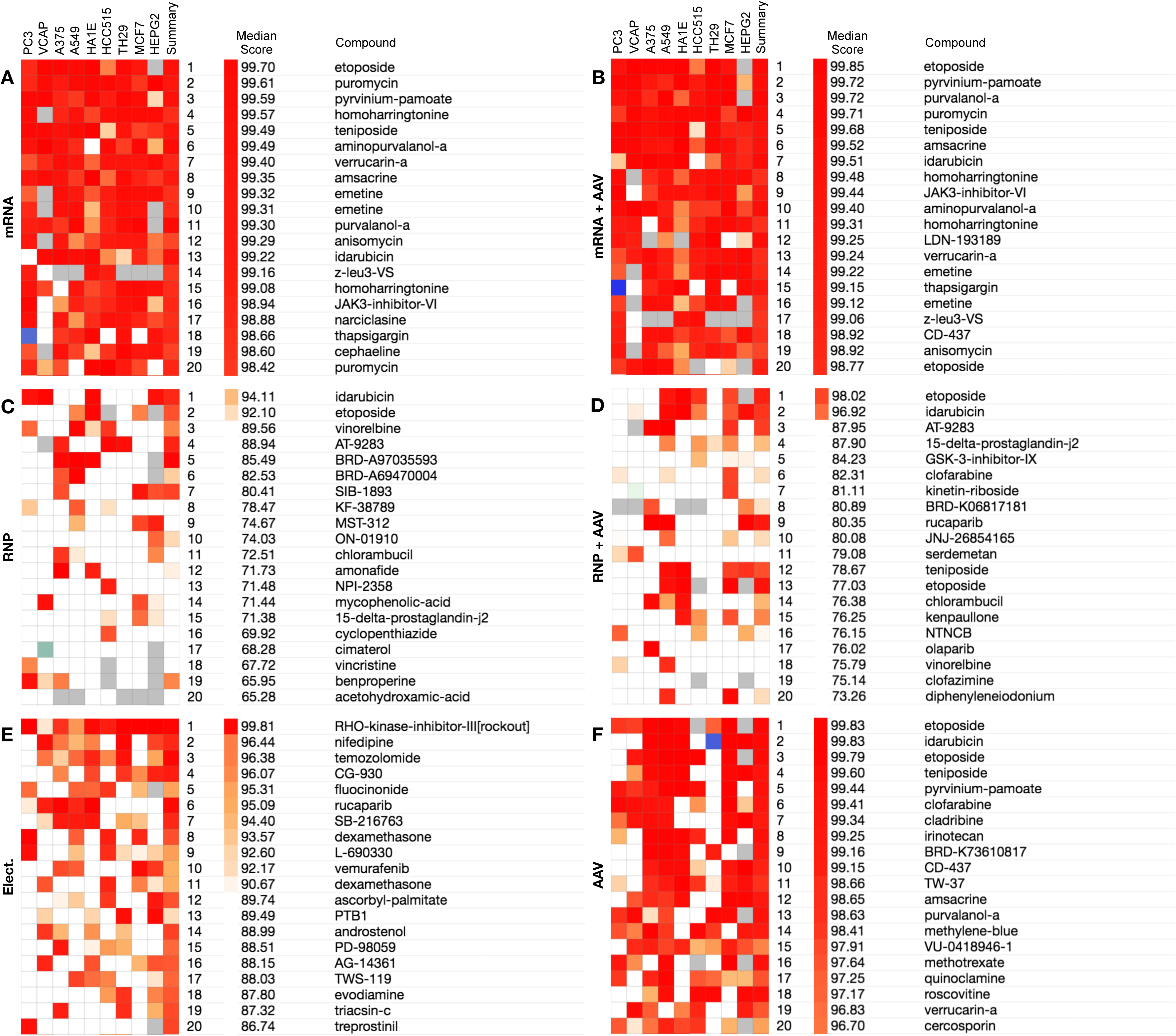
Small molecules with a positive connectivity across various cell lines in CMap to different components of the CRISPR/Cas9-AAV6 based genome editing machinery: (**A**) mRNA, (**B**) mRNA + AAV, (**C**) RNP, (**D**) RNP + AAV, (**E**) Mock and (**F**) AAV. The summary/ median score in each case refers to the median of connectivity scores (including the Summary score) across the nine cell lines in CMap-PC3, VCAP, A375, A549, HA1E, HCC515, TH29, MCF7 and HEPG2. Methods detailed in Subramanian *et al.*

**Figure 2:**
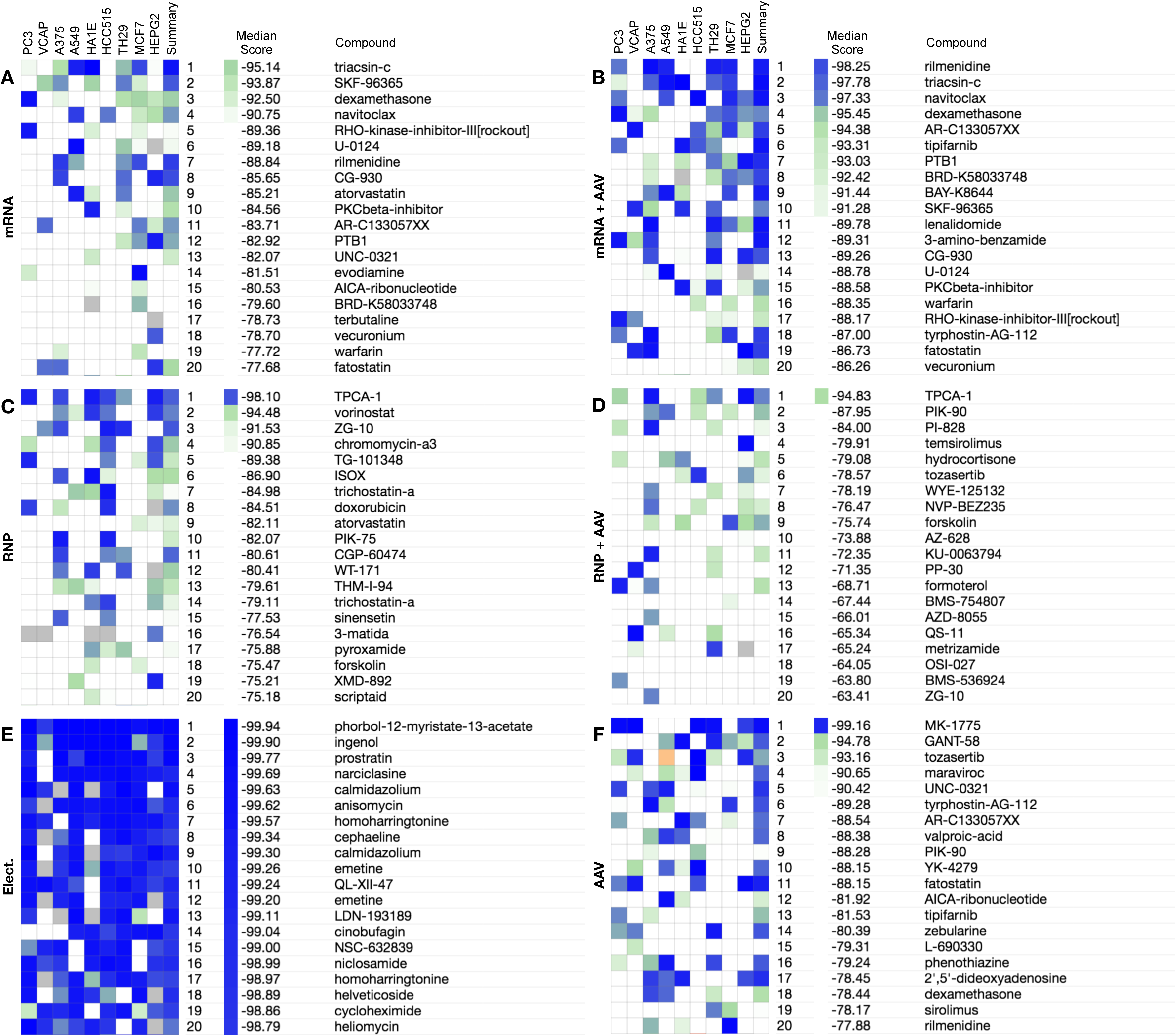
Small molecules with a negative connectivity across various cell lines in CMap to components of CRISPR/Cas9-AAV6 based genome editing machinery: (**A**) mRNA, (**B**) mRNA + AAV, (**C**) RNP, (**D**) RNP + AAV, (**E**) Mock and (**F**) AAV. The median score in each case refers to the median of connectivity scores (including the Summary score) across the nine cell lines in Cmap. Methods detailed in Subramanian *et al.*

Next, we considered small molecule compounds in CMap with a negative connectivity score to query signatures for each of the genome editing components, that is, suggesting dissimilarity or negative correlation between the small molecules and transcriptional responses to CRISPR/Cas9-AAV6 components (Figure 2 and Supplementary Table S2). Among the top 20 small molecules showing negative connectivity score to Cas9 mRNA, three – triacsin-C, rilmenidine and terbutaline – are adrenergic receptor agonists. Furthermore, adrenergic receptor agonists also appeared in the top 20 small molecules with high negative connectivity to the other components of the editing machinery (Figure 2B to 2F): triascin-C for mRNA + AAV; rilmenidine for AAV and mRNA + AAV; and formoterol for RNP + AAV. Previously, a screen of approximately 4,000 small molecules [12] identified and confirmed two small molecules – brefeldin A and L755507, an adrenergic receptor agonist – as capable of improving CRISPR-mediated homology directed repair (HDR), further supporting potential associations between adrenergic receptors agonists and CRISPR/Cas9 gene editing. Interestingly, we also found that 7 out of the top 20 small molecules with negative connectivity to RNP were histone deacetylase inhibitors (HDACi) – namely, vorinostat, ISOX, trichostatin-A, WT-171, THM-I-94, pyroxamide and scriptaid. In addition, we found that transcriptional signatures of other epigenome modifying drugs or small molecules including the HDACi valproic acid are also in opposite direction (negatively correlated) to that of AAV. Separately, we found that the histone lysine methyltransferase inhibitor UNC-0321 shows negative connectivity to Cas9 mRNA or AAV and the DNA methyltransferase inhibitor zebularine has negative connectivity to AAV (Supplementary Table S2). Consistent with these observations, trichostatin was recently shown to enhance CRISPR/Cas9 enabled targeted nucleotide substitutions [13]. We provide more information on the top 20 molecules with negative connective score to each of the components in Supplementary Table S2.

In interpreting the significance of our findings, it is important to note a number of factors and limitations. First, there is no single parameter for assessing outcomes or the efficiency of CRISPR/Cas9 gene editing as it is dependent on several factors including: rates of transfection of the editing components into cells, entry of editing components into nuclei, on- and off-target cleavage events, repair of cleaved sites, survival of edited cells and ultimately phenotypic traits associated with the target genomic locus. Transcriptional responses to genome editing components may be related to any of these factors as well as to other unknown factors. Our results suggest that small molecules with shared transcriptional responses to the various gene editing components may influence distinct aspects of these processes. For example, some of the small molecules identified by our approach have been shown to influence CRISPR/Cas9 editing efficiency through their effects on distinct DNA repair processes: while trichostatin – an HDACi – enhances targeted nucleotide substitutions in CRISPR/Cas9 edited cells [13], L755507 enhances HDR [12]. Thus, small molecules could offer a means to fine-tune specific cellular processes associated with gene editing technologies.

Second, up-regulation or down-regulation of genes by CRISPR/Cas9-AAV6 may involve a subset three subsets of genes: (*i*) those that have no effect on genome editing outcomes; (*ii*) those that have positive effect on editing; and (*iii*) those that have a negative effect on editing outcomes. Small molecules that affect expression of genes whose expression has no effect on editing outcomes would be expected to similarly have no or little effect on editing outcomes, those that up-regulate genes whose up-regulation has a positive effect on editing would potentiate or be synergistic to editing while those that up-regulate genes that have a negative effect on editing would have a negative effect on editing. In contrast, small molecules that down-regulate genes that have a positive effect on editing would be more likely to have a negative effect on editing outcomes while those that down-regulate genes that have a negative effect on editing would have a positive effect on eventual editing outcomes. Thus, applying our observations to enhance or suppress genome editing outcomes would require further experimentation to identify the impact of transcriptional effects of the small molecules on eventual editing outcomes at the phenotypic level.

Third, our study uses CMap data that is based on transcriptional perturbations of cancer cell lines none of which includes the CD34+ stem cell that was used to obtain transcriptional effects of CRISPR/Cas9-AAV6 by Cromer *et al*. Hence, some of the responses may be cell line specific. Furthermore, even the same cancer cell line can harbor extensive clonal diversity that could cause variability in responses to the same perturbations [14]. We address this limitation by considering responses across multiple cancer cell lines which increases the likelihood the responses may be conserved across several cell types.

Finally, some of the small molecules considered in this study are experimental and others include approved drugs. This opens the possibility of interactions between genome editing technologies and drugs. Such interactions may occur *ex vivo*, for example, when attempting to edit patient derived cells obtained from a patient on a medication that has transcriptional effects interfering with editing components. In a few cases, small molecule interactions with genome editing components may also occur *in vivo*, albeit less likely in the clinical context with a few exceptions whereby editing may be done *in vivo*. For example, treatment of retinal dystrophies may involve *in vivo* genome editing [15] which could expose the editing machinery to interactions with medications in a patient. In the laboratory setting, whole animals such as a mouse line with a molecular barcode generating CRISPR/Cas9 system integrated in the germline and active in somatic cells [16] may be exposed to small molecules as part of unrelated experiments thereby potentially influencing any tracing of individual cells based on the CRISPR/Cas9 generated barcodes. Interactions between CRISPR/Cas9 and small molecules could also be an important factor to consider in functional genomic screens for drug mechanism of action where CRISPR/Cas9 is used to knock-out genes [17, 18]. For example, in such cases if a small molecule interferes with CRISPR/Cas9 editing, the drug may obscure or enhance the ability to knockout its target by CRISPR/Cas9.

While previous studies have suggested a number of small molecules that could modulate CRISPR/Cas9 gene editing, the results presented here suggest that additional small molecules with a similar effect may be predicted using transcriptional data. In conclusion, transcriptional responses to genome editing components could potentially be used to predict small molecules that modulate CRISPR/Cas9 editing and may be applicable in a combination, hybrid therapeutic approach in which small molecules are administered concomitantly with gene editing machinery.

## ACKNOWLEDGEMENTS

This work was supported by the Association pour la recherche en cancerologie de Saint-Cloud (ARCS), ADEBIOPHARM, and the OpenHealth Institute. We thank Dominique Bellet for helpful discussions and feedback.

## AUTHORS’ CONTRIBUTIONS

A.V.N. and G.H.S. conceived and directed the study. A.V.N. and G.H.S. analyzed and interpreted the data with the support of M.J.A. A.V.N. and G.H.S. wrote the manuscript. M.J.A. revised the manuscript. All authors read and approved the final manuscript.

---

**Supplementary Table S1:** Top 20 small molecules with the highest positive connectivity to each component of the CRISPR/Cas9 genome editing machinery. Mechanism of action for each of the small molecules is provided.

**Supplementary Table S2:** Top 20 small molecules with the highest positive connectivity to each component of the CRISPR/Cas9 genome editing machinery. Mechanism of action for each of the small molecules is provided.

